# Personalised whole-body modelling links gut microbiota to metabolic perturbations in Alzheimer’s disease

**DOI:** 10.1101/2025.10.28.685084

**Authors:** Tim Hensen, Lora Khatib, Lucas Patel, Daniel McDonald, Antonio González, Siamak MahmoudianDehkordi, Colette Blach, Alzheimer Gut Microbiome Project Consortium, Rob Knight, Rima Kaddurah-Daouk, Ines Thiele

## Abstract

The human gut microbiome has been linked to metabolic disturbances in Alzheimer’s disease (AD). However, the mechanisms by which gut microbes might influence metabolic dysfunction in AD remain poorly understood. Previously, we used constraint-based metabolic modelling to associate an increased risk of AD with altered production of microbiome-derived metabolites. In this study, we investigated whether these previous results can also be identified in AD patients. Therefore, we created personalised whole-body metabolic models from gut metagenomics samples from 34 AD dementia patients, 51 individuals with mild cognitive impairments, and 298 healthy controls. These *in silico* models were profiled to predict the metabolic influences of gut microbiomes on blood metabolites with previously reported alterations in AD. We found an increased capacity of AD host-microbiome co-metabolism to produce S-adenosyl-L-methionine, L-arginine, creatine, taurine, and formate in the blood of AD dementia patients and patients with mild cognitive impairments. The metabolic predictions were then mechanistically linked to gut microbial changes in AD. This analysis identified that increased relative abundances of *Bacteroides uniformis* and *Bacteroides thetaiotamicron* were key factors driving the predicted metabolic changes. Furthermore, the predicted altered microbial influences on blood metabolites were also associated with allelic variations in the *APOE* risk gene in healthy individuals, which confirmed our previous findings. In conclusion, we identified blood metabolites whose perturbations in AD may be influenced by gut microbiota and predicted the key microbial drivers for these metabolic influences. These findings may facilitate the development of microbiome-informed treatments of AD.

## 1 Introduction

Alzheimer’s disease (AD) is the most common neurodegenerative disease. AD is generally characterised by a progressive loss of neurons, extracellular accumulation of amyloid beta proteins, and intracellular accumulation of hyperphosphorylated tau proteins^1,2^. In addition, several major risk factors for AD have been identified, such as old age, lower cognitive functioning, and being a carrier of the *APOE ε*4 allele^3^. However, the mechanisms linking these risk factors to AD remain poorly understood. Emerging evidence has shown that microbiota inhabiting the gastrointestinal tract may influence AD pathology by acting on the gut-brain axis, which comprises bidirectional communication pathways connecting gut microbes to the brain^4^. Previous reports have linked several microbial metabolites to AD, such as secondary bile acids^5^, short-chain fatty acids^6^, indoles^7^, and trimethylamine N-oxide^8^. However, the network-wide effects of microbial contributions to the gut-brain axis in AD remain largely unknown.

In addition to alterations in microbial metabolites, metabolomic studies have identified widespread metabolic perturbations in AD patients. A recent systematic review by Yin et al.^9^ identified 396 AD-associated metabolites across 66 pathways. Commonly found perturbed pathways in AD were arginine and proline metabolism, glutamate metabolism, phenylalanine metabolism, cysteine and methionine metabolism, and taurine and hypotaurine metabolism^9^. To elicit the effects of gut microbiota on these perturbed pathways and identify the metabolic perturbations influenced by the gut microbiome, computational modelling methods have enabled the prediction of isolated gut microbiome influences on human metabolism^10,11^, independent of other drivers of metabolism, such as disease progression, dietary habits, and environmental exposures.

*In silico* genome-scale metabolic models of gut microbiome communities have been a powerful tool for predicting the metabolic host-microbiome crosstalk in health and disease^10,12^. These metabolic microbiome community models integrate metagenomic data with comprehensive microbial genome-scale metabolic reconstruction resources, such as AGORA2^13^ and APOLLO^14^, and enable mechanistic investigations of personalised gut microbiome metabolism using constraint-based optimisation methods, such as flux balance analysis (FBA)^15^. Microbiome community models have been previously used to predict altered metabolic features of gut microbiomes in inflammatory bowel diseases^16,17^, colorectal cancer^18^, and Parkinson’s disease^19,20^. Microbiome community models, however, cannot model the bidirectional metabolic influences that the host and gut microbiome exert on each other, nor can they investigate microbial influences on human metabolites. To model host-microbiome crosstalk, recent studies have incorporated personalised microbiome community models into sex-specific and organ-resolved whole-body metabolic models (WBMs)^21^, creating integrated models of host-microbiome co-metabolism. These microbiome-personalised WBMs have been previously utilised to study host-microbiome co-metabolism on inflammatory bowel diseases^22^ and AD dementia^23,24^.

Previously, we have utilised gut microbiome-personalised WBMs to predict the contributions of gut microbiota to blood metabolites that may influence AD phenotype, and could associate these predictions with major risk factors of AD, including increasing age, *APOE ε*4 carrier status, and global cognitive scores^24^. This prior analysis associated the predicted gut microbiome influences on arginine metabolism, methionine metabolism, and secondary bile acid metabolism with an increased risk of developing AD, and identified *Eggerthella lenta* as a potential key contributor to the secondary bile acids, deoxycholate and lithocholate. However, our previous study had important limitations. The predicted metabolic associations with the AD risk factors were made for healthy ageing individuals and have not been validated in an independent cohort of AD patients. Secondly, poor correlations have been observed between the metabolic predictions and plasma metabolomics from the same samples. This lack of data agreement has been attributed to previously reported challenges that may arise when comparing human gut metagenomic and plasma metabolomic abundances, such as incomplete metagenomic and metabolomic annotations, temporal mismatches in sample collection, and differential interactions with factors, such as host genetics, dietary intake, environmental interactions, and disease progression^25,26^. However, the use of lower-resolution 16S rRNA metagenomic data may have also reduced the modelling capacity to predict plasma abundances in corresponding samples. This previous study also identified potential microbial contributors to those metabolic predictions. However, the methods used in that study could not identify microbial contributors to non-microbial metabolites with microbe-derived precursors. Secondly, this prior study could not account for mechanistic constraints on microbial contributions to circulatory metabolites that may arise from model-specific diet–microbe–host interactions.

In this study, we followed up on our prior predictions by testing and validating these findings in an independent cohort that included individuals with mild cognitive impairment (MCI) and AD dementia patients. We also assessed whether the previously observed lack of associations between metabolic predictions and plasma metabolomics may be improved with high-resolution shotgun metagenomics data and expanded genome-scale reconstruction libraries for metagenomic mapping. Furthermore, we introduced a novel approach for identifying the microbial contributors to metabolic predictions in microbiome-personalised WBMs, which accounted for indirect microbial contributions on non-microbial metabolites and the context-specific contribution potentials in each microbiome-personalised WBM. Finally, this study replicated our previous results in healthy individuals by associating the metabolic predictions with *APOE ε*4 carrier status and Montreal Cognitive Assessment (MoCA) scores. Our analysis revealed that the gut metagenomes of MCI and AD dementia patients had increased capacities to influence circulatory metabolic levels of S-adenosyl-L-methionine, L-arginine, creatine, taurine, and formate, confirming and expanding our previous predictions. Our results also confirmed that *in silico* host-microbiome predictions on circulatory metabolites and plasma metabolomic abundances may not be straightforwardly correlated. Our approach for identifying key microbial contributors to human metabolites determined that increased relative abundances of *Bacteroides uniformis* and *Bacteroides thetaiotaomicron* were a key factor for the predicted AD-related metabolic shifts in blood. Lastly, we demonstrated that the predicted circulatory metabolic shifts in MCI and AD dementia patients were also associated with *APOE* allele carrier status in healthy individuals. Taken together, this study identified progressive shifts in gut microbial contributions to circulatory metabolites from healthy to MCI and AD dementia phenotypes, and showed that *APOE ε*4 carrier may be involved in preclinical host-microbiome crosstalk.

## 2 Results

We aimed to predict the metabolic influences of gut microbiota on a curated list of 38 blood metabolites in MCI and AD patients. To that end, we obtained gut metagenomics samples from a cohort of 383 American participants that included 298 healthy controls, 51 MCI patients with a presumptive AD diagnosis, and 34 AD dementia patients. The gut metagenomic samples were used to generate microbiome-personalised WBMs, which were given an average Western metabolic diet. FBA^15^ was then performed to interrogate the models on their maximal flux capacity to accumulate each metabolite in the WBM blood compartment (Figure 1, Methods). The predicted metabolic fluxes were associated with MCI and AD status to identify potential changes in metabolic contributions from gut microbiota. We then tested whether the *in silico* flux predictions could be correlated with plasma metabolite abundances in a subset of 168 individuals. Next, we identified the key microbial contributors to metabolites of interest by first performing feature selection to reduce the search space of possible microbe-metabolite associations and then determining the best associations between the flux predictions and associated microbial species. Lastly, we validated our prior findings on AD risk markers by associating the AD-associated flux predictions with MoCA scores and *APOE* genotype status in a healthy subcohort.

**Figure 1:**
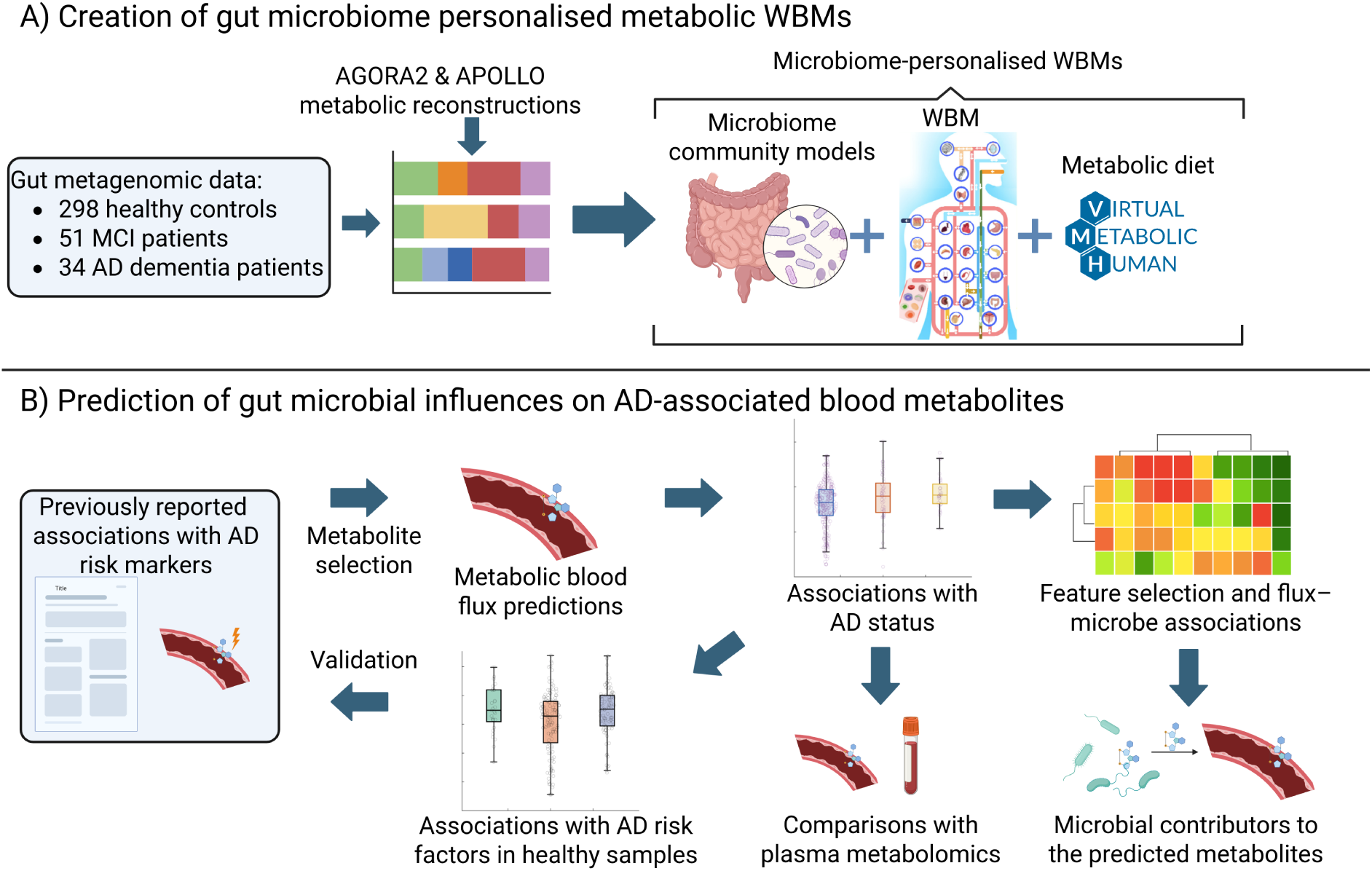
Schematic overview of the study design. A) Metabolic reconstructions of microbial species in the AGORA2^13^ and APOLLO^14^ resources were mapped onto the microbial species in the gut metagenomics samples. The mapped microbial reconstructions were then combined into microbiome community models and personalised with their corresponding relative abundances. These microbiome community models were then used to personalise standard male and female WBMs corresponding to the sex of each microbiome sample donor, and were given an average Western metabolic diet. B) Metabolic blood fluxes were then predicted for 38 previously analysed blood metabolites with reported metabolomic changes in AD patients. The flux predictions were associated with MCI and AD dementia patients to find altered gut microbial influences on these metabolites in MCI and AD patients. The predicted metabolic associations were then compared with corresponding plasma metabolomic abundances from the same individuals. Next, the microbial contributors to the predicted fluxes were identified by applying a novel feature selection approach and then determining the best microbe-metabolite associations. Lastly, the predicted metabolites with altered flux predictions in MCI or AD dementia patients were associated with Montreal Cognitive Assessment (MoCA) scores and *APOE* genotype in a subset of healthy individuals to validate prior WBM modelling predictions. This figure was created in BioRender.

### 2.1 Study cohort

In this study, we used gut metagenomic data from a cross-sectional cohort of 400 participants from a pooled sample of ten Alzheimer’s Disease Research Centres that were part of the Alzheimer Gut Microbiome Project (Methods). The cohort included 308 cognitively healthy controls, 53 individuals with MCI and a presumptive diagnosis of AD, and 39 individuals with diagnosed AD dementia. However, after removing samples from individuals with co-morbid diseases and samples with low metagenomic sample quality (Methods), 383 metagenomic samples remained available for analysis, including 298 healthy controls, 51 MCI patients with a presumptive diagnosis of AD, and 34 AD dementia patients (Table 1). The healthy individuals in the analysed cohort were on average 73.29 years of age (Standard deviation, SD=7.53), while the MCI and AD dementia patients were 76.33 years (SD=7.45) and 76.65 years (SD=6.69) of age, respectively. Furthermore, women were over-represented in the healthy control group (67.11% female samples), but not in the MCI (49% female samples) and dementia (41% female samples) groups. Other available metadata in the cohort were the body mass index (BMI), MoCA scores, years of education, and *APOE* genotype. A lower BMI was observed in AD dementia patients compared to both the cognitively healthy and MCI participants. Furthermore, no difference in the years of education was observed between healthy, MCI, and AD dementia participants. The MoCA test scores showed a strong, consistent decline in cognitive abilities with dementia progression. Finally, a higher proportion of *APOE ε*4 positive individuals was observed in the AD dementia patients (38.24%) compared to cognitively healthy individuals (29.19%). This increased proportion was not significant after testing for over-representation of *APOE ε*4 carrier status (Methods, Table 1).

**Table 1:**
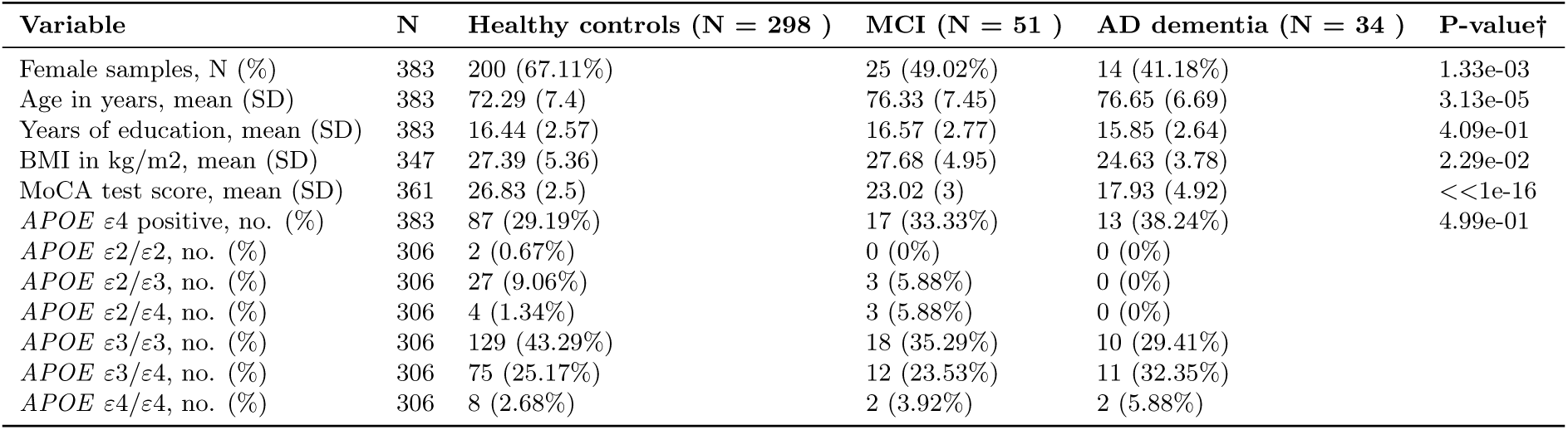
Cohort characteristics of healthy controls, AD patients with mild cognitive impairment (MCI), and AD dementia patients. †P-values were calculated based on one-way ANOVA tests and with *χ*^2^ tests of independence for all binary variables.

| Variable | N | Healthy controls (N = 298 ) | MCI (N = 51 ) | AD dementia (N = 34 ) | P-value† |
| --- | --- | --- | --- | --- | --- |
| Female samples, N (%) | 383 | 200 (67.11%) | 25 (49.02%) | 14 (41.18%) | 1.33e-03 |
| Age in years, mean (SD) | 383 | 72.29 (7.4) | 76.33 (7.45) | 76.65 (6.69) | 3.13e-05 |
| Years of education, mean (SD) | 383 | 16.44 (2.57) | 16.57 (2.77) | 15.85 (2.64) | 4.09e-01 |
| BMI in kg/m2, mean (SD) | 347 | 27.39 (5.36) | 27.68 (4.95) | 24.63 (3.78) | 2.29e-02 |
| MoCA test score, mean (SD) | 361 | 26.83 (2.5) | 23.02 (3) | 17.93 (4.92) | <<1e-16 |
| *APOE* ε4 positive, no. (%) | 383 | 87 (29.19%) | 17 (33.33%) | 13 (38.24%) | 4.99e-01 |
| *APOE* ε2/ε2, no. (%) | 306 | 2 (0.67%) | 0 (0%) | 0 (0%) | |
| *APOE* ε2/ε3, no. (%) | 306 | 27 (9.06%) | 3 (5.88%) | 0 (0%) | |
| *APOE* ε2/ε4, no. (%) | 306 | 4 (1.34%) | 3 (5.88%) | 0 (0%) | |
| *APOE* ε3/ε3, no. (%) | 306 | 129 (43.29%) | 18 (35.29%) | 10 (29.41%) | |
| *APOE* ε3/ε4, no. (%) | 306 | 75 (25.17%) | 12 (23.53%) | 11 (32.35%) | |
| *APOE* ε4/ε4, no. (%) | 306 | 8 (2.68%) | 2 (3.92%) | 2 (5.88%) | |

### 2.2 Generation of gut microbiome-personalised WBMs

To create the microbiome-personalised WBMs, we first mapped the species-level taxonomies in the gut metagenomics data onto microbial genome-scale metabolic reconstructions in the AGORA2^13^ and APOLLO resources^14^, which contained 7,302 human microbial strains and 247,092 genome-specific metabolic reconstructions of human microbes, respectively. After taxonomic mapping, we renormalised the relative abundances and removed three additional microbial species that were below the relative abundances cut-off of one in a million in at least one sample. This procedure mapped a total of 430 microbial species onto AGORA2 and APOLLO, which corresponded to an average read coverage of 71% (SD=16%) compared to the pre-mapped metagenomic reads (Table 2, Table S1).

**Table 2:** The effects of taxonomic mapping on the microbiome read counts and diversity metrics. The before-and-after mapping columns display the mean and standard deviation of read counts and diversity statistics before and after taxonomic mapping on the AGORA2 and APOLLO resources. The fold changes indicate the mean relative change after taxonomic mapping for the statistics. Fold change values below one indicate a reduction in read counts and ecological diversity after taxonomic mapping. The Shannon and Simpson indices are diversity metrics that take into account both the species richness and ecological evenness in a sample. The Shannon index is more influenced by species richness, while the Simpson diversity index is more influenced by ecological evenness. The lower diversity indices after mapping indicate that the taxonomic diversity is reduced as a result of the mapping procedure. The mean Shannon fold change is lower than the mean Simpson fold change, indicating that species richness might be more affected than ecological evenness. †P-values were calculated using a two-sample t-test.

| Statistic | Before mapping | After mapping | Fold change | P-value† |
| --- | --- | --- | --- | --- |
| Mean read counts (SD) | 6,993,476 (2,706,890) | 5,043,143 (2,431,164) | 0.714 (0.16) | <<1e-16 |
| Total number of named microbial species | 1,299 | 430 | 0.33 |  |
| Mean species richness (SD) | 149.48 (45.59) | 61.272 (19.733) | 0.46 (0.235) | <<1e-16 |
| Mean Shannon index (SD) | 3.101 (0.5) | 2.529 (0.463) | 0.843 (0.235) | <<1e-16 |
| Mean Simpson diversity index (SD) | 0.892 (0.08) | 0.841 (0.106) | 0.953 (0.169) | 7.99e-14 |

Taxonomic mapping further reduced species richness from an average of 149.48 (standard deviation, SD=45.59) to 61.27 (SD=19.73, Table 2). The Shannon and Simpson diversity indices also showed a loss in microbial diversity after mapping. Notably, the relative decrease in the Simpson diversity index was less severe than the relative reduction in Shannon indices. This observation suggests that the ecological evenness could be less affected than the loss in taxonomic richness, as the Simpson diversity index is more influenced by ecological evenness than the Shannon index^27^. Furthermore, taxonomic mapping changed the relative abundances of the most abundant phyla, *Bacteroides* and *Firmicutes*, from 50.05% (SD=19.44%) to 64.78% (SD=20.36%) and from 35.86% (SD=17.33%) to 21.46% (SD=15.51%), respectively (Table S1). The mapping procedure also increased the relative abundances of the most abundant microbial species, *B. uniformis*, which changed from an average of 10.23% (SD=12.15%) before mapping to 13.48% (SD=14.99%) after mapping. After taxonomic mapping, the filtered metagenomic reads were used to create personalised microbiome community models, which were joined with the large intestinal lumen of sex-matched male and female WBMs (Methods). The generated microbiome-personalised WBMs (Table S2) were also given a diet corresponding to the metabolic intake of an average Western dietary intake^28^ (Table S3). In summary, the microbiome-personalised WBMs captured on average 71.4% (SD = 16%) of the reads in the metagenomic data.

### 2.3 Selection of metabolites for *in silico* modelling

To predict microbial influences on metabolic alterations in AD patients, we curated a set of 38 microbially produced metabolites that were analysed in the previous WBM modelling studies on microbial links to AD risk factors^24^ and AD status^23^. The selected metabolites (Methods, Table S4) included twelve biochemical subclasses, of which “bile acids, alcohols, and derivatives” (18 metabolites) was the most abundant. Other abundant biochemical subclasses were “amino acids and peptides” (six metabolites) and benzenediols (four metabolites). All 38 selected metabolites were present in the WBM blood compartments, of which 22 could be produced by gut microbes in the WBMs. The remaining sixteen metabolites had metabolic precursors with microbial origins. Furthermore, eleven of the analysed metabolites were also present in the defined Western diet, while twelve metabolites could only be produced by gut microbial species (Table S4). Although the selected set of metabolites likely captured only a small part of all microbially influenced metabolites in AD patients, the analysed metabolites were chosen to achieve our specific aims of validating previously predicted gut microbiome-driven metabolic associations with AD or risk factors of AD and for assessing the validity of negative findings in these previous modelling results.

### 2.4 Prediction of gut microbiome influences on AD-related metabolites in blood

Previously, we have shown that gut microbiome influences on L-arginine, S-adenosyl-L-methionine, deoxycholate, and lithocholate were associated with increased risk for AD^24^. To test whether these prior findings may persist in MCI and AD dementia, we predicted the maximal metabolic flux for the accumulation of the selected metabolites in the blood of the microbiome-personalised WBMs. After processing the predicted fluxes, we performed a multivariate outlier inspection (Methods) and identified two outlier samples, which were subsequently removed. We then associated the predicted fluxes with cognitive status by performing logistic regressions on the flux predictions. Control variables were added for sex, age at stool collection, and relevant technical covariates (see Methods). The regressions were performed in a pairwise manner, i.e., healthy versus MCI, MCI versus dementia, and healthy versus dementia. Five metabolites were identified to potentially associate with MCI or AD dementia status. These metabolites were S-adenosyl-L-methionine, L-arginine, creatine, taurine, and formate (Figure 2, Table S5). All five metabolites had higher predicted flux values in MCI or AD dementia compared to healthy controls. No changes between MCI and AD dementia patients were found in the flux predictions. However, this lack of change between MCI and AD dementia patients might be explained by all the analysed MCI patients also having a presumptive AD diagnosis. Nonetheless, the median values and regression estimates of the predicted fluxes were consistently higher in AD dementia patients compared to MCI patients, potentially indicating a consistent change with AD status. No significant associations with MCI or AD dementia status were detected for secondary bile acids. However, we did observe weakly increased predicted fluxes from healthy controls to MCI for deoxycholate (p=0.08) and lithocholate (p=0.09, Table S5). No such association was present for deoxycholate and lithocholate when comparing healthy controls and AD dementia patients. Taken together, we predicted increased potential gut microbiome contributions to S-adenosyl-L-methionine, L-arginine, creatine, taurine, and formate in blood. These results align with our prior findings and identify creatine and formate as circulatory metabolites with altered gut microbial influences in AD. Importantly, as these findings result from a targeted analysis, they are unlikely to represent the full spectrum of AD-associated gut microbiome influences on blood metabolites.

**Figure 2:**
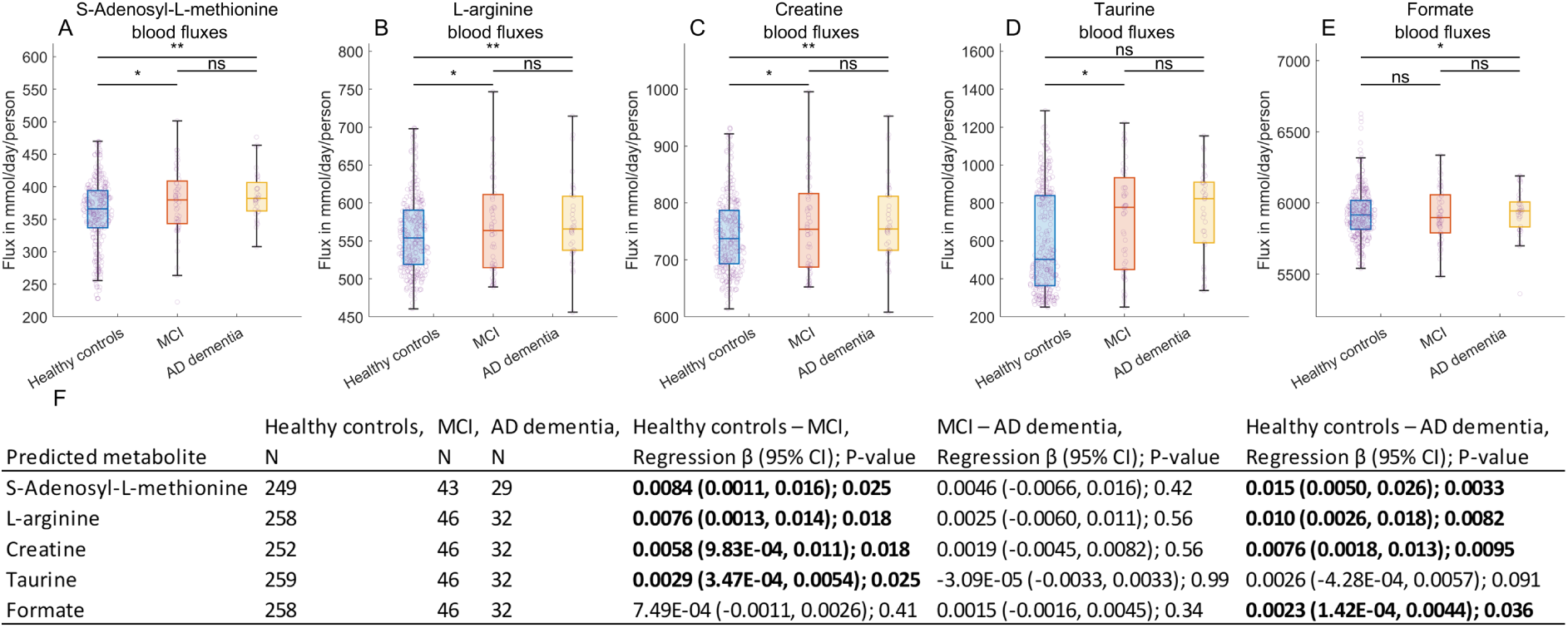
Predicted host-microbiome production capacities for circulatory metabolites. **A-E.** shows the predicted fluxes in the blood of healthy controls (blue), MCI (red), and AD dementia patients (yellow) for the analysed metabolites that were associated with either MCI or AD status. The asterisks above each box plot indicate the significance level. Two asterisks indicate that P*<*0.01, one asterisk means that P*<*0.05, while “ns” indicates that no association was identified. **F.** Shows the regression table summarising the logistic regression results for the shown metabolites in A-E. The number of samples (N) for which fluxes could be predicted is shown for healthy controls, MCI, and AD dementia patients. For each regression, the regression *β*, 95% CI, and regression p-value are shown. The bold text indicates that the obtained P value was below 0.05. The regression *β* shows the log odds ratios for the change in probability of a more severe AD state when increasing the blood flux values. The positive *β* values indicate that higher predicted blood fluxes associate with more severe cognitive decline.

### 2.5 The predicted blood fluxes did not associate with plasma abundances from the same samples

We assessed whether the predicted metabolic fluxes would correlate with plasma metabolomic abundances from a subset of 168 individuals in the analysed cohort, including 130 healthy controls, 28 MCI patients, and 10 AD dementia patients. L-arginine, creatine, and taurine have been measured in the plasma samples, but not S-adenosyl-L-methionine and formate. To evaluate whether the flux predictions were predictive of the plasma abundances, linear regressions were performed (Methods) on the predicted L-arginine, creatine, and taurine fluxes against their corresponding plasma abundances. In line with previous observations^23,24^, no association was found for the three tested metabolites (Table S6). We then assessed whether the relative plasma metabolomic abundances of L-arginine, creatine, or taurine associated with AD progression, as previously reported^29–34^. However, no association with MCI or AD status was found for these metabolites in these samples (Table S6). This lack of signal in the metabolomic data may be explained by the lower sample numbers for the plasma metabolomics, as only 28 MCI and 10 AD dementia patients were included. Taken together, this analysis did not observe correlations between the *in silico* flux predictions and corresponding plasma metabolomics, concurring with previous observations^23,24^. This result may further indicate that genome-scale modelling predictions on gut microbiome contributions to circulatory metabolites cannot be straightforwardly compared to corresponding metabolomic data from blood samples.

### 2.6 Identification of microbial contributors to blood fluxes associated with AD status

Next, we identified the microbial species that influenced the altered flux predictions in MCI and AD dementia patients. These flux-associated microbial species were identified by 1) utilising the mechanistic structure of the WBMs to find the set of microbial species that influenced the predicted metabolites, and 2) by using regularised regressions to filter on the most sparse set of microbial species that were most predictive of the fluxes (Methods). To identify which microbial species could have contributed to the predicted flux values, we analysed the FBA shadow price solutions for the microbial species biomass compounds in each microbiome-personalised WBM. These shadow prices measured the sensitivity of the predicted fluxes to a hypothetical change in the abundance of a microbial species. For each predicted metabolite, the mean microbial species biomass shadow price values across all analysed samples were calculated, along with the associated 95% confidence intervals (CI) around the mean (Methods). Microbial species were then removed from the list of potential microbial influencers if their mean shadow price value was not statistically different from zero, i.e., the 95% CI around the mean microbial species biomass shadow price included zero. Out of the total 430 microbial species in the microbiomes, 303 potential microbial influencers of S-adenosyl-L-methionine were identified. For L-arginine and creatine, 286 and 289 microbial influencers were found, respectively, while for taurine and formate, 325 and 304 potential microbial influencers were found (Table 3, Table S7).

**Table 3:** Summary statistics on the number of microbial species that limited the blood flux predictions for the selected metabolites. Columns two to four show the total number of potential microbial contributors, the mean, and the standard deviation of microbial contributors across the cohort. Columns five to seven show the same summary statistics for a filtered set of analysed microbial contributors to the predicted fluxes. The last column shows the explained variance of the analysed microbial species for the associated flux predictions. The explained variance was calculated after fitting elastic net regression models (see Methods) with the flux predictions as the response variable and predictors for the analysed microbial species and with sex as the control variable.

| Metabolite | Identified microbial contributors |  |  | Analysed microbial contributors |  |  |  |
| --- | --- | --- | --- | --- | --- | --- | --- |
| | Total | Mean | SD | Total | Mean | SD | Explained variance ($R^2$) |
| S-adenosyl-L-methionine | 303 | 58.1 | 19.5 | 8 | 4.37 | 1.81 | 0.799 |
| L-arginine | 286 | 51.3 | 16.3 | 15 | 6.32 | 2.84 | 0.996 |
| Creatine | 289 | 54.8 | 17.2 | 16 | 6.83 | 3.11 | 0.996 |
| Taurine | 325 | 61.6 | 20.2 | 11 | 5.63 | 2.67 | 0.999 |
| Formate | 304 | 56.4 | 17.9 | 20 | 9.37 | 4.00 | 0.803 |

However, these microbial species could not have equally contributed to the flux predictions due to differences in their sample presence and relative abundances. To identify the most important microbial influencers of the predicted fluxes, we performed feature selection using the stability selection method^35^, which applies LASSO regressions on bootstrapped samples^36^ to select the minimal set of microbial species that were most predictive of the flux predictions (Methods). This step drastically narrowed the list of flux-associated microbial species (Table 3, Table S7). For S-adenosyl-L-methionine, eight key microbial species were identified, which, after controlling for sex, explained 79.9% of variance in the S-adenosyl-L-methionine flux predictions (Methods). L-arginine and creatine had fifteen and sixteen flux-associated microbes, respectively, that could explain 99.6% of the variance in both the L-arginine and creatine flux predictions. For taurine and formate, eleven and twenty key microbial species were identified that explained 99.9% and 80.3% of their associated fluxes after controlling for sex, respectively. To summarise, we utilised the microbe-associated shadow prices and a multivariate feature selection method to find potential key microbial species for each of the five metabolites with AD-associated fluxes. The identified sets of flux-associated microbial species were highly predictive of the flux results, confirming that the identified microbial sets contained major drivers of the flux predictions.

### 2.7 *B. uniformis* and *B. thetaiotaomicron* were key drivers for the increased metabolic flux predictions with AD patients

After identifying the set of microbial influencers of the flux predictions, we aimed to identify the key microbes driving the AD-associated flux predictions. To find these microbial species, we performed elastic net regressions^37^ (Methods), a multivariate method for identifying relevant features in omics datasets^38^. The elastic net regressions were performed on the z-scaled flux predictions with covariates for the selected flux-associated microbes and sex as the control variable. These regressions identified *B. uniformis* as the most important microbial contributor to the AD-associated predicted metabolites, except for formate, which was highly influenced by *Escherichia coli* (Figure 3, Table S8).

**Figure 3:**
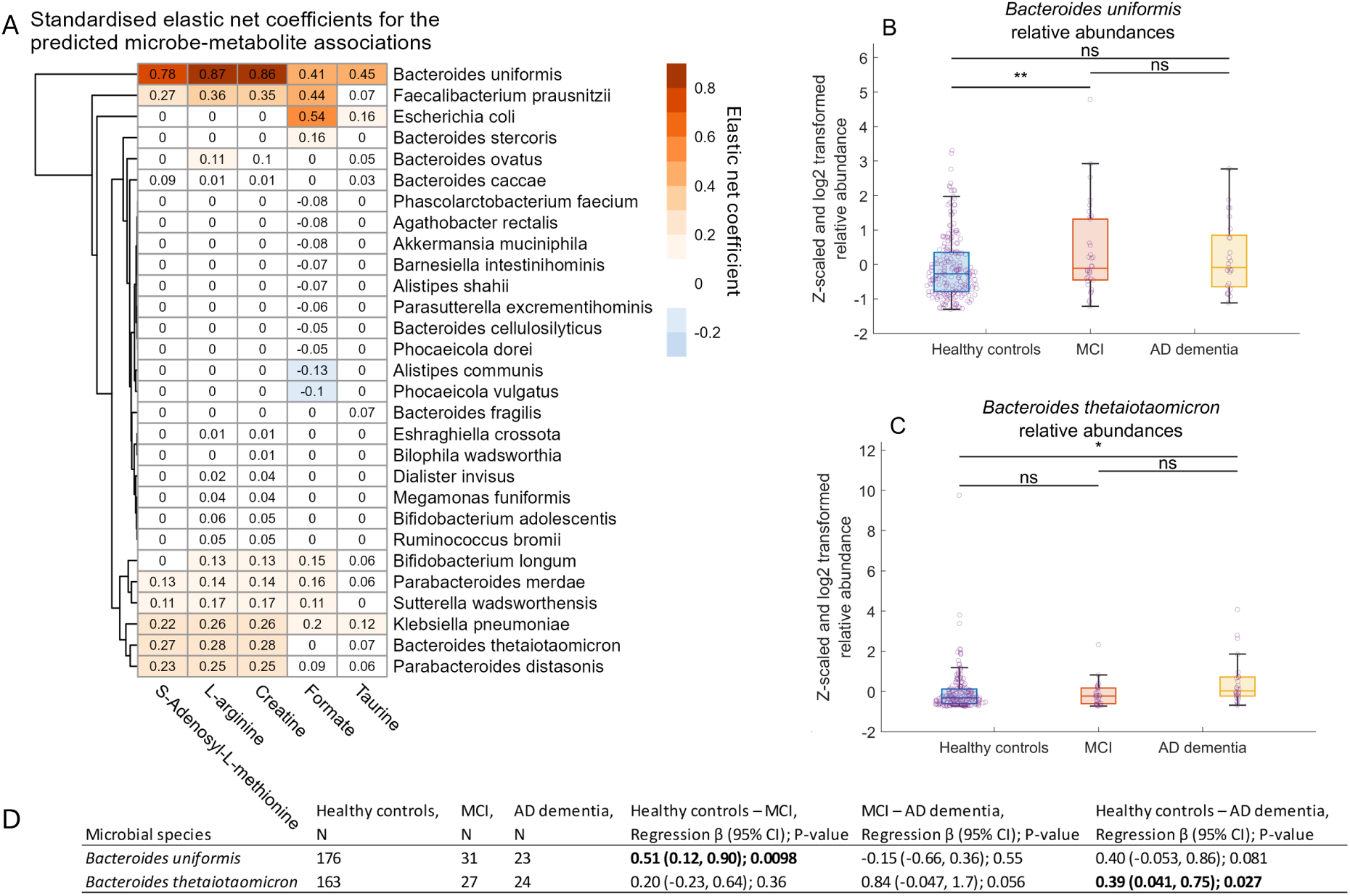
**A.** Standardised elastic net coefficients for the selected microbial contributors to the predicted metabolites. The standardised coefficients represent the predicted change in flux value in standard deviations resulting from a one-standard-deviation change in standardised microbial relative abundance. High positive coefficients (red) indicate a strong positive association between blood metabolic flux predictions and microbial species relative abundances. Conversely, negative coefficients (blue) suggest negative flux-microbe associations. Coefficients of zero indicate that a microbial species was not identified to be associated with the corresponding flux prediction. The elastic net coefficients show that *B. uniformis* is a key microbial contributor to the flux predictions of all analysed metabolites. **B** and **C**. Box plots of microbial relative abundances of *B. uniformis* and *B. thetaiomicron* in healthy controls, MCI, and AD dementia patients. Both microbial species had higher relative abundances in AD patients compared to controls. No other flux-associated microbial species was associated with MCI or AD status. The asterisks above each box plot indicate the significance level. Two asterisks indicate that P*<*0.01, one asterisk means that P*<*0.05, while “ns” indicates that no association was identified. **D**. Regression table summarising the logistic regression results of the shown microbial species in B and C. The number of samples (N) for which fluxes could be predicted is shown for controls, MCI patients, and dementia patients. For each regression, the regression *β*, 95% CI, and regression p-value are shown. The bold text indicates that the obtained P value was below 0.05. The regression *β* shows the log odds ratios for the change in probability of a more severe AD state when increasing the blood flux values. The positive *β* values show that higher relative abundances of *B. uniformis* and *B. thetaiomicron* were associated with AD dementia.

*E. coli* was also associated with the predicted taurine blood fluxes, but did not associate with S-adenosyl-L-methionine, L-arginine, and creatine. Another major contributor to the predicted blood fluxes was *Faecalibacterium prausnitzii*, which was the second most important influencer of the flux predictions for all metabolites, except for taurine. Furthermore, we found *B. thetaiotaomicron* to be a key contributor to the predicted creatine, L-arginine, and S-adenosyl-L-methionine fluxes. Finally, the elastic net models found a negative coefficient for *Alistipes communis* with the formate flux predictions, indicating that increased growth of *A. communis* was associated with lower formate flux predictions. Next, we tested whether these microbial keystones were also associated with the AD stages by performing the same logistic analysis as described for the flux predictions (Methods). These logistic regressions found increased relative abundances of *B. uniformis* and *B. thetaiotaomicron* in AD dementia patients (Figure 3B-D, Table S8), suggesting that these microbes could have driven the increased flux predictions in MCI patients and AD dementia patients. Taken together, we identified the key microbial species for influencing the AD-associated flux predictions and found increased relative abundances of the key microbes, *B. uniformis* and *B. thetaiotaomicron*, in MCI patients and AD dementia patients.

### 2.8 The predicted AD-associated blood fluxes were also associated with *APOE ε*2 and *APOE ε*4 carrier status in healthy individuals

Next, we tested whether the five identified AD-associated metabolites would also have different predicted fluxes in healthy individuals with lower or higher risk for developing AD. Therefore, we used only the 298 gut metagenomic samples from cognitively healthy individuals and investigated differences in the flux predictions in allelic risk groups for the *APOE* gene. There are three allelic variations of the *APOE* gene, *ε*2, which protects against AD pathogenesis, *ε*3, which is the normal risk allele, and *ε*4, which is linked with increased risk for AD^39,40^. These three alleles give rise to six allelic combinations (*ε*2/*ε*2, *ε*3/*ε*3, *ε*4/*ε*4, *ε*2/*ε*3, *ε*2/*ε*4, and *ε*3/*ε*4). As the *ε*2/*ε*2 and *ε*4/4 allelic combinations had low abundances in our samples (Table 1), we combined the individuals with the *ε*2/*ε*2 and *ε*2/*ε*3 genotypes into the *APOE ε*2 group. Individuals with *ε*3/4 and *ε*4/4 were conversely grouped into the high-risk *ε*4 group. The normal risk *ε*3 group included only the *APOE ε*3/*ε*3 genotype. Individuals with the *APOE ε*2/*ε*4 genotype were excluded from this analysis due to the simultaneous expression and potential counteracting effects on microbiota of both the neuroprotective *APOE ε*2 and the AD-risk associated *APOE ε*4 alleles. ANCOVA regressions were performed on pairwise associations between two *APOE* groups (predictor), i.e., *ε*2 versus *ε*3, *ε*3 versus *ε*4, and *ε*2 versus *ε*4, and the blood flux predictions for the 38 analysed metabolites (Methods). The regressions found associations with the *APOE* risk groups for all analysed metabolites except for creatine, which narrowly missed the significance threshold (Figure 4, Table S9).

**Figure 4:**
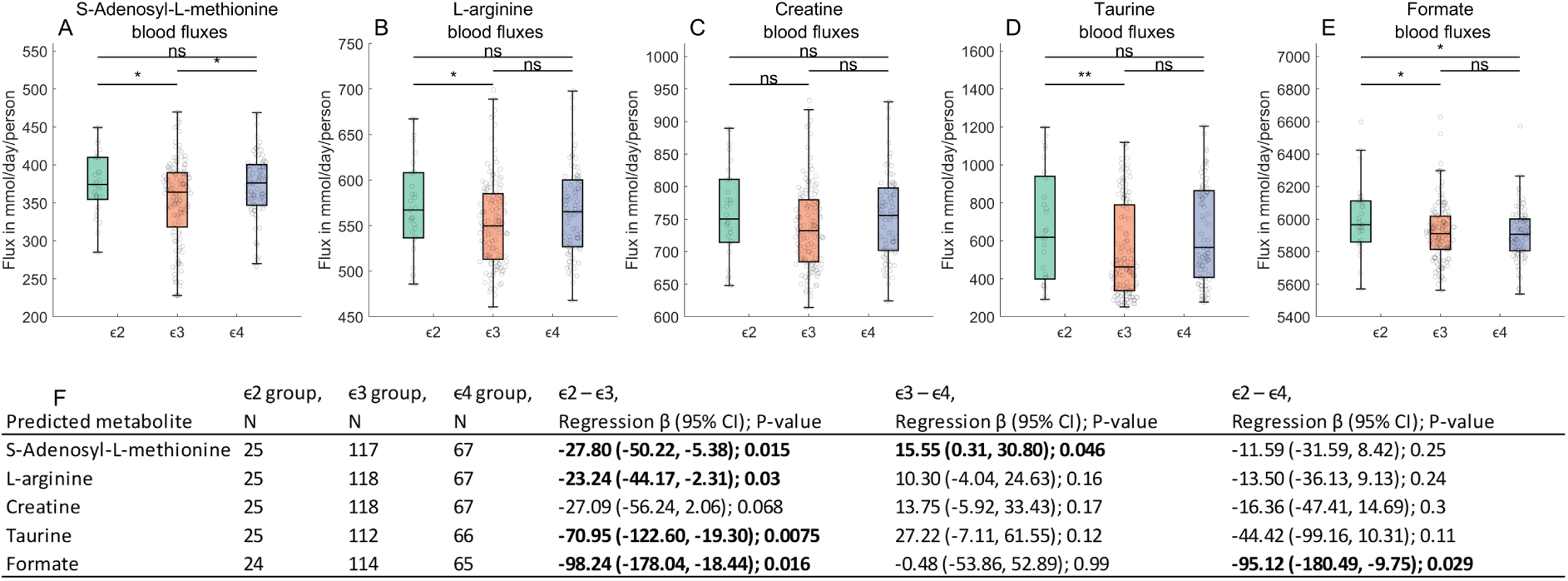
Predicted host-microbiome blood production capacities in healthy individuals for the selected circulatory metabolites. **A-E**. shows the predicted blood fluxes for the *ε*2 (green, *ε*2/2 & *ε*2/3), *ε*3 (orange, *ε*3/3), and *ε*4 (blue, *ε*3/4 & *ε*4/4) *APOE* risk groups. The purple dots indicate the predicted flux values in each group. The asterisks above each box plot indicate the significance level. Two asterisks indicate that P*<*0.01, one asterisk means that P*<*0.05, while ns indicates that no association was identified. **F.** Regression tables for the metabolites shown in A-E. Results are shown for each pairwise group comparison, i.e, *ε*2 versus *ε*3, *ε*3 versus *ε*4, and *ε*2 versus *ε*4. The results were obtained through pairwise ANCOVA regressions on the *APOE* risk groups (predictor of interest) against the flux predictions (response), while controlling for household and relevant technical covariates (Methods). The *β* and 95% confidence intervals (CI) show the average change in flux value from the first group, e.g., *ε*2, to the second group, e.g., *ε*3. The bold text indicates that the obtained P value was below 0.05. These results show lower blood flux predictions in *ε*3 for taurine, L-arginine, and S-adenosyl-L-methionine compared to both *ε*2 and *ε*4, and lower blood fluxes in the *ε*4 group for formate compared to the *ε*2 group.

When comparing the low-risk *ε*2 group to the high-risk *ε*4 group, differential fluxes were only found for formate, which had lower fluxes in the *ε*4 group. When comparing the low-risk *ε*2 group to the normal-risk *ε*3 group, higher fluxes were found in the *ε*2 group for S-adenosyl-L-methionine, L-arginine, and taurine, potentially indicating similar bacterial influencers of these metabolites. Finally, higher S-adenosyl-L-methionine flux predictions were found in the *ε*4 group compared to the *ε*3 group. We then assessed whether the previously identified key microbial drivers of the AD-associated flux predictions may have also driven the differential fluxes with the *APOE* groups. Therefore, we performed additional regressions to associate their relative abundances with *APOE*, as described for the flux predictions. These regressions could, however, not find any differences in relative abundances between the *APOE* risk groups for *B. uniformis* and *B. thetaiotaomicron*, suggesting that the differential fluxes across the *APOE* groups might not have been driven by these microbial species. We further investigated whether the analysed metabolites had altered fluxes in healthy individuals with differing MoCA scores. MoCA is a popular psychometric test with high sensitivity for detecting subjective and mild cognitive impairments^41^. To analyse potential associations, we performed linear regressions on the predicted blood fluxes from healthy individuals (predictor) with the MoCA test scores as the response variable (Methods). However, none of the analysed metabolites correlated with MoCA test scores in the plasma metabolomics (Table S9). To summarise, the analysed AD-associated flux predictions differed with *APOE* risk groups in healthy individuals from the analysed cohort, but not with the MoCA test scores.

## 3 Discussion

In this study, we examined the metabolic influences of gut microbiota on circulatory metabolites that may influence the AD phenotype. To that end, we predicted the host-microbiome production capacities for blood metabolites in microbiome-personalised WBMs in healthy controls, MCI, and AD dementia. Our main results were that i) previously predicted microbiome-driven metabolic associations in at-risk healthy individuals for S-adenosyl-L-methionine and L-arginine were replicated in MCI and AD dementia. ii) We further confirmed previous observations that predicted blood fluxes may not be straightforwardly correlated with corresponding metabolomic plasma measurements. iii) We introduced a novel approach for identifying the key microbial contributors to predicted circulatory metabolites and determined that *B. uniformis* was the key contributor to the AD-associated blood flux predictions. iv) We showed that the predicted AD-associated host-microbiome metabolite production capacities were also associated with *APOE* genotype in healthy individuals.

Our analysis revealed that the predicted gut microbiome contributions to S-adenosyl-L-methionine, L-arginine, creatine, taurine, and formate in blood increased in MCI and AD compared to healthy controls. However, no changes were observed between MCI and AD dementia patients. This result aligns with previous systematic reviews by Ticinesi et. al (2018)^42^ and by Jemimah et. al (2022)^43^, who suggested that changes in gut dysbiosis may not always be identified between MCI and AD dementia. The predicted increased microbial contributions to S-adenosyl-L-methionine may suggest a gut microbial influence on DNA methylation, as S-adenosyl-L-methionine is the main cofactor for DNA methylation^44^. This result aligns with previous results from post-mortem brain samples, which identified increased S-adenosyl-L-methionine abundances in AD patients^45,46^. However, lower cerebrospinal fluid levels of S-adenosyl-L-methionine have also been associated with a higher risk for AD progression in MCI patients^47^. Lower S-adenosyl-L-methionine abundances have been previously proposed to result from vitamin B12 deficiency^48^, which is associated with AD incidence^49^. S-adenosyl-L-methionine is mainly sourced in the liver from L-methionine and ATP^50^. Our result suggests that the gut microbiome may influence methionine metabolism by increasing the availability of its precursors. However, it remains difficult to ascertain how these microbial influences could alter plasma levels of S-adenosyl-L-methionine, as our models do not account for the complex regulatory feedback loops of methionine metabolism^51^. The predicted increase in gut microbiome contributions to L-arginine in AD patients could have diverse effects on human metabolism, as L-arginine is a key precursor for several metabolites that associate with AD pathology, such as nitric oxide, L-glutamate, GABA, polyamines, and creatine^9^. L-arginine has been proposed to potentially mitigate cognitive decline by reducing oxidative stress^52^. However, the effects of microbial influences on L-arginine metabolism may be diverse, as increased L-arginine abundances have also been associated with AD in post-mortem brain tissues^53,54^, while lower L-arginine abundances in plasma have been linked with higher cognitive function in old age^55^. Moreover, multiple gut microbiome-driven contributions may interact to produce deleterious effects. For example, S-adenosyl-L-methionine can provide the methylation substrate for the production of asymmetric dimethylarginine from L-arginine. Dimethylarginine is a known antagonist of nitric oxide production and has been previously linked to Alzheimer’s disease^56^ and cardiovascular dysfunction^57,58^, which itself is a major risk factor for Alzheimer’s disease^59^. The increased S-adenosyl-L-methionine and L-arginine fluxes might also link with the predicted increased microbial influences of taurine and L-arginine. S-adenosyl-L-methionine is a precursor of taurine via the production of S-adenosyl-homocysteine, which is reduced to cysteine, which is in turn oxidised and decarboxylised to produce taurine^60,61^. Conversely, creatine biosynthesis occurs through the production of guanidinoacetate from L-arginine and glycine, which is then methylated by S-adenosyl-L-methionine to produce creatine^62^. Both taurine and creatine have important functions in human health. Creatine is the major source for phosphocreatine, which is the key cellular storage molecule for ATP^63^. Taurine has functions in human physiology that range from bile acid detoxification^64^ to osmoregulation^65^ and blood pressure regulation^66^. However, taurine has also been reported to protect against neurodegeneration by refolding misfolded proteins^67^ and can potentially mitigate amyloid *β* accumulation^68,69^. Finally, our results included increased predicted microbial influences on formate. This result may suggest a microbial link with its direct precursor, formaldehyde, which is a neurotoxic compound and known antagonist of methylation^70^. However, lower formate urine and faecal levels have also been reported in AD patients^23,71^, suggesting a more complex and nuanced role of formate production in AD patients.

In our prior study, we associated L-arginine, S-adenosyl-L-methionine, deoxycholate and lithocholate with increased risk for AD in healthy individuals. We validated these prior predictions by showing that L-arginine and S-adenosyl-L-methionine were also associated with MCI and AD. Although no significant signal was identified for deoxycholate and lithocholate blood fluxes, we did observe a tendential increase in the flux predictions for these metabolites in AD patients (Table S5), which is in line with the mounting evidence for increased microbial conversion of bile acids in AD progression from MCI^5,72,73^. Taken together, our targeted analysis confirmed previous observations and expanded the set of circulatory metabolites, which may be differentially influenced by gut microbiota in AD.

Previous whole-body modelling studies were unable to successfully correlate *in silico* flux predictions with plasma^24^ or urine^23^ metabolomics. Similarly, we did not identify significant associations between predicted fluxes and plasma metabolomic data. This result suggests that distal metabolomic readouts, such as those obtained from blood or urine, may not be readily comparable to flux predictions derived from microbiome-personalised WBMs. This interpretation aligns with previous reports showing that direct correlations between gut metagenomic data and plasma metabolomics are challenging to establish^25,26,74^. One plausible explanation for the absence of flux–plasma associations in this study is that the five identified metabolites are either common dietary compounds or predominantly synthesised *de novo* in humans. For example, creatine and taurine can be obtained from animal-source foods^75^, meaning that plasma alterations may reflect dietary differences between healthy individuals and AD patients. Circulatory S-adenosyl-L-methionine is furthermore primarily sourced from hepatic synthesis^50^, whereas L-arginine levels may also be driven by increased protein catabolism or altered endogenous synthesis^76^. Similarly, circulating formate concentrations could be explained by altered *de novo* synthesis from the folate cycle^77^. Because the plasma abundances of these metabolites can be explained by non-microbial sources, shifts in gut microbiome contributions may not directly translate into measurable plasma changes. Nonetheless, microbial contributions could still exert effects on disease-associated molecular pathways and thereby influence AD progression.

We also identified the key microbial contributors to the five metabolites with altered blood fluxes in AD. To achieve this result, we developed a two-step feature selection strategy that utilised the mechanistic structure of microbiome-WBMs and statistical feature selection to reduce the search space of possible microbe-metabolite associations. This approach was highly effective in identifying the most predictive microbial contributors to the flux predictions, while remaining mechanistically grounded and accounting for the modelled dynamics of host–microbiome co-metabolism. Our analysis revealed that *B. uniformis* and *B. thetaiomicron* were major drivers of the increased flux predictions in MCI and AD dementia. *B. uniformis* is a commensal gut bacterium with reported probiotic properties^78,79^. Similarly, higher relative abundances of *B. thetaiomicron* were identified as strong contributors to the elevated flux predictions in AD patients. This species is also a well-known commensal with potential probiotic effects, including reducing oxidative stress^80^ and alleviating ulcerative colitis symptoms^81^. In addition, *E. coli* was identified as a major contributor to formate flux predictions (Figure 3). Although *E. coli* itself was not associated with AD status, it has been implicated in AD pathology due to its ability to produce amyloid-*β* peptides, such as curli fibres, which promote amyloid-*β* aggregation^82^. Taken together, our feature selection strategy identified *B. uniformis* and *B. thetaiomicron* as key microbial contributors to the five analysed circulatory metabolites. While their roles in AD-related metabolic perturbations require further validation, these findings implicate both species as potential microbial targets for future therapeutic strategies.

Lastly, we observed that the metabolites with altered microbial contribution potentials in MCI and AD dementia were also associated with *APOE* genotype in healthy individuals. Whether the allelic variations of the *APOE* gene exert causal effects on the microbiome or whether the associations are purely correlational remains uncertain. Nevertheless, growing evidence supports a link between *APOE* genotype and gut microbial composition^83–87^. Notably, our observed associations with *APOE* carrier status replicate our previous findings on S-adenosyl-L-methionine in healthy individuals^24^, reinforcing a possible connection between *APOE* genotype and gut microbiome interactions with human methionine metabolism. More broadly, these findings are consistent with the hypothesis that gut microbial dysbiosis may arise before the onset of measurable cognitive decline^88^, which would make the gut microbiome a potential early marker of dementia, as demonstrated previously^89^

### 3.1 Strengths and limitations

The main strength of this study lies in the use of unique metabolic modelling methods, the richly annotated shotgun metagenomics dataset, and the availability of associated plasma metabolomics for investigating the metabolic effects of gut microbiota in AD patients. The combined mapping on both the AGORA2^13^ and APOLLO^14^ resources further ensured a high read coverage of the gut microbiomes in the metabolic models, enabling more accurate investigations of host-microbiome co-metabolism in AD patients. One limitation of the study was the lack of data on dietary habits, meaning that diet differences between AD patients and healthy controls could have confounded the results. Another limitation was the limited number of metabolomic samples that overlapped with the metagenomics data, especially for AD patients. Furthermore, the modelling predictions were limited by inherent features of the COBRA framework, such as the assumption of steady state metabolism, which does not capture the dynamics of host-microbe and microbe–microbe interactions. In addition, the models could not capture the physiological changes of gut dysbiosis in AD patients, such as increased intestinal permeability^90^. Previously, physiologically based pharmacokinetic models have successfully modelled gut microbiome-mediated intestinal metabolic absorption^91^ and have been integrated with WBMs^92^. Future studies might integrate physiologically based pharmacokinetic models with microbiome-personalised WBMs to improve modelling predictions on gut microbial contributions to metabolites in the bloodstream.

### 3.2 Conclusions

In conclusion, we identified shifts in potential gut microbiome contributions to circulatory metabolites that may influence AD pathology. We also determined that these predicted metabolic shifts may be explained by increased relative abundances of *B. uniformis* and *B. thetaiomicron* using a novel feature selection and microbe-metabolite association approach. We further showed that AD-associated microbial contributions to the analysed predicted metabolites were also associated with *APOE* genotype in healthy individuals, suggesting a potential link between *APOE*, the gut microbiome, and AD pathogenesis. Although experimental validation will be necessary to confirm these predicted microbe-metabolite associations in AD, we hope that these results will help guide and inspire future efforts to uncover the key mechanisms of host-microbiome co-metabolism in AD dementia.

## 4 Materials and methods

### 4.1 Participant recruitment

Participants were recruited from ten multiple NIA-funded Alzheimer’s Disease Research Centres that were part of the Alzheimer Gut Microbiome Project (AGMP), including UC San Diego, UC Davis, Stanford University, Kansas University, Wisconsin University, Indiana University, New York University, Wake Forest University, Cleveland Clinic, Nevada, and the University of Alabama at Birmingham. Before admission into the ADRC, studies, participants provided written informed consent following the Declaration of Helsinki. After admission, participants underwent neuropsychological testing according to the standardised UDS 3.0 framework, which included the MoCA assessment. *APOE* genotypes were obtained from the whole-blood samples using competitive allele-specific polymerase chain reaction (PCR)–based KASP genotyping assays (LGC Genomics, Beverly, MA)^93^. Presumptive etiologic diagnoses of MCI due to AD and dementia due to AD were made by a multidisciplinary diagnostic panel, and were made based on the diagnostic criteria from the National Institute on Aging–Alzheimer’s Association^94,95^.

### 4.2 Metabolomics

Plasma samples were collected by the NCRAD fluid biomarker program on behalf of the AGMP. The plasma metabolomics data were generated from EDTA plasma samples by Metabolon Inc. (Durham, NC, USA). The raw plasma dataset included 1,665 biochemicals, 1,316 compounds of known identity (named biochemicals) and 349 compounds of unknown structural identity (unnamed biochemicals). After plasma sample collection, the samples were immediately stored at -80 degrees Celsius, after which they were prepared using the MicroLab STAR® system from the Hamilton Company. During the sample preparation, small molecules attached to proteins were recovered by first dissociating the small molecules bound to the proteins and then precipitating the proteins with methanol by shaking for 2 minutes (Glen Mills GenoGrinder 2000) and centrifugation. The extracts were then divided into five different aliquots. One aliquot for each of four analysis methods and one aliquot as a backup. Then, organic solvents were removed by placing the samples on a TurboVap® (Zymark), after which they were stored overnight under nitrogen.

The dried sample aliquots were all analysed using Waters ACQUITY ultra-performance liquid chromatography (UPLC) and a Thermo Scientific Q-Exactive high resolution/accurate mass spectrometer interfaced with a heated electrospray ionisation (HESI-II) source and Orbitrap mass analyser operated at 35,000 mass resolution. After overnight storage, the sample extracts were reconstituted in solvents compatible with each of their associated analysis methods. One sample aliquot was analysed in acidic positive ion conditions optimised for more hydrophilic compounds (PosEarly). The extract from this aliquot was gradient eluted from a C18 column (Waters UPLC BEH C18-2.1x100 mm, 1.7 µm) using water and methanol, containing 0.05% perfluoropentanoic acid (PFPA) and 0.1% formic acid (FA). Then, a second aliquot was analysed using the same acidic positive ion conditions, but optimised for more hydrophobic compounds (PosLate). Gradient elution was performed from the same C18 column using methanol, acetonitrile, water, 0.05% PFPA and 0.01% FA, but was operated at an overall higher organic content. A third aliquot was analysed using basic negative ion optimised conditions and a separate dedicated C18 column (Neg). The basic extracts of this aliquot were gradient eluted from the column using methanol and water, however, with 6.5mM Ammonium Bicarbonate at pH 8. The fourth aliquot was analysed using negative ionisation following elution from a HILIC column (Waters UPLC BEH Amide 2.1x150 mm, 1.7 µm) in a gradient consisting of water and acetonitrile with 10mM Ammonium Formate, pH 10.8 (HILIC). The MS analysis was performed by alternating between MS and data-dependent MSn scans using dynamic exclusion. The scan range covered 70-1000 m/z. The raw data were then extracted, peak-identified, and QC-processed using Metabolon’s hardware and software. Peak quantification was performed by calculating the area under the curve. A data normalisation step was further performed to correct for variation from instrument inter-day tuning differences. In this step, each compound was corrected by registering the medians to equal one and normalising each data point proportionately in run-day blocks.

After obtaining the corrected metabolomic peaks, metabolites with over 25% missing data points were excluded from the analysis. To account for sample variability, probabilistic quotient normalisation was applied, followed by a log2 transformation. The remaining missing values were then imputed using the k-nearest neighbour algorithm. Outlier samples were identified and removed using the local outlier factor method implemented in the bigutilsr R package. To address extreme single concentration values, any measurement with an absolute abundance exceeding q = abs(qnorm(0.0125/n)), where n is the number of samples, was marked as missing. This threshold q corresponds to values with less than a 2.5% two-tailed probability of originating from the same normal distribution as the other measurements, incorporating a Bonferroni-like correction (by dividing by the sample size). The newly marked missing values were then imputed in a second round using the k-nearest neighbour algorithm. Batch effects were corrected using the COMBAT package.

### 4.3 Gut metagenomics

The analysed faecal samples were collected between March 2022 and June 2023 as previously described^96^. Briefly, the samples were collected at home using standardised faecal collection kits. Collected samples were returned the same day via overnight delivery in insulated containers that were chilled with frozen gel packs. Upon return, the samples were weighed, scored on the Bristol stool scale, and stored at -80 degrees Celsius until further processing.

Sample DNA extraction was performed using the MoBio PowerMag Soil DNA isolation kit with a magnetic bead plate. The extracted genomic DNA (gDNA) was quantified using the Quant-iT PicoGreen double-stranded DNA (dsDNA) assay kit from Thermo Fisher Scientific Inc. The samples then underwent miniaturised KAPA HyperPlus library preparation using the iTru indexing strategy^97,98^. The prepared libraries were normalised, pooled based on their concentrations, and PCR-cleaned. Size selection was performed based on the 300-700 bp range using the Sage Science PippinHT pipeline. DNA sequencing was then performed on an Illumina NovaSeq 6000 as a paired-end 150-cycle run at the University of California, San Diego (UCSD) IGM Genomics Centre.

The generated sequence data were then processed, trimmed, and filtered using the qp-fastp-minimap2 pipeline version 2023.12^99,100^, as described previously^101^. The reads were filtered using minimap2 against the GRCh38.p13, T2T-CHM13v2.0, viral Phi X 174, and the Human Pangenome Reference assemblies^102^. The processed metagenomic sequencing data were then uploaded to Qiita^103^ (Study ID: 15448) for further processing. Taxonomic classification was performed using the Woltka pipeline^104^ version v0.1.4 (qp-woltka 2023.11), which mapped the metagenomic data against the Web of Life 2 reference database. Lastly, the SCRuB^105^ pipeline was used to statistically correct the processed read table for well-specific microbial contamination and well-to-well sample leakage across the 96-well plate.

### 4.4 Taxonomic mapping

The processed gut metagenomics data were mapped onto pan-species genome-scale metabolic reconstructions in the combined AGORA2^13^ and APOLLO^14^ resources, using the MARS pipeline^106^. After the mapping procedure, the metagenomic reads were normalised by calculating the relative read abundances per sample. To limit false positives in the mapped microbiomes, sample relative abundances below the threshold of 0.00001% were removed, and the remaining read abundances were re-normalised. The mapping coverages, i.e., the fraction of available metagenomic data in a sample after mapping, were calculated from the original and mapped relative abundances after removing reads with relative abundances below the threshold. After taxonomic mapping, an additional quality control step was performed, which resulted in samples being removed if they had fewer than 1 million reads after taxonomic mapping or a mapping read coverage below 25%.

### 4.5 Microbial metabolic reconstructions

Genome-scale metabolic reconstructions for the mapped microbial species were retrieved from the AGORA2 (version 2.01, https://www.vmh.life/files/reconstructions/AGORA2/) and APOLLO^14^ (version 1.0, https://doi.org/10.7910/DVN/PIZCBI) resources. As the metabolic reconstructions in AGORA2 and APOLLO are strain-resolved, pan-species models were generated using the *createPanModels.m* function^107^. The created pan-species models represented the combined metabolic content of their associated strain-resolved reconstructions and contained the union set of all reactions, metabolites, and genes in the species-associated strain-level reconstructions. Furthermore, the pan-species models contain a new pan-species biomass reaction that was created by taking the union set of consumed metabolites in the associated strain-level reconstructions. For consumed metabolites present in more than one biomass reaction in the associated strain-level reconstructions, the mean stoichiometric coefficient was calculated for the consumed metabolite in the pan-species biomass reaction.

### 4.6 Microbiome community models

Personalised metabolic models of gut microbiome communities were created using the mgPipe module in the Microbiome Modelling toolbox 2.0^108^. In mgPipe, personalised microbiome community models are created by joining the mapped pan-species models from a sample in a shared microbiota lumen and adding transport reactions to enable metabolic exchanges between the pan-species models and the shared microbiota lumen. Then, a dietary compartment and faecal compartment are connected to the microbiota lumen compartment to enable dietary intake and faecal excretion of microbial products. Next, the mapped relative abundances were integrated into the microbiome communities via the addition of a community biomass reaction (VMH reaction ID: communityBiomass), which lists each microbe’s biomass reaction with the microbe’s relative abundance as the stoichiometric coefficient. This community biomass reaction ensured proportional microbial growth for each integrated pan-species model per the mapped relative species abundances. Consequently, the community biomass reaction ensured that the metabolic contribution capacity of each microbial species to the host was proportional to their relative abundances.

### 4.7 Whole-body metabolic models

We used the generic male and female WBMs^21^ (version 1.04c^109^) for the host component of the microbiome-personalised WBMs. These organ-resolved WBMs captured the metabolism of 26 organs and 13 biofluid compartments, which were connected in an anatomically accurate manner. The male generic WBM contained 83,395 reactions, 58,095 metabolites, and 105,936 constraints, while the female generic WBM contained 85,892 reactions, 60,537 metabolites, and 109,757 constraints. All parameters were set to their defaults^21^.

### 4.8 Microbiome-personalised WBMs

The microbiome-personalised WBMs were created using the *combineHarveyMicrobiota.m* function, which integrates the personalised microbiome community models with the large intestinal lumen compartment of the generic WBMs, as described previously^21^. This function also introduces transport reactions to enable metabolic exchanges between the microbiota and the host. This function also updated the coupling constraints for each microbe’s reaction to their biomass reactions, to a ratio of 1:400, following previous WBM modelling studies^21,23,24,110^.

Further parameterisation of the WBMs was performed by setting the upper and lower flux bounds of the *Excretion EX microbiota LI biomass[fe]* reaction to one mmol/day/person. This reaction excretes the *microbiota LI biomass* compound, which is produced by the *communityBiomass* reaction. Fixing the flux bounds of the *Excretion EX microbiota LI biomass[fe]* reaction ensured that all generated WBMs produced the same metabolic flux over their microbiome community, regardless of differences in community compositions. In addition, the upper and lower flux bounds of the *Whole body objective rxn* were set to one mmol/day/person, following previous WBM studies^21,23,24,111^. The *Whole body objective rxn* includes a linear combination of the whole-body organ biomass productions and associated coefficients for the proportional organ weights. This additional constraint ensured proportional organ biomass production between samples, corresponding to the organ weights.

### 4.9 Diet

The microbiome-personalised WBMs were further parameterised with dietary uptake reactions and reaction constraints from the Virtual Metabolic Human database (https://www.vmh.life)^28^. Reactions and constraints were chosen for the average Western diet, since sufficient information on dietary uptake was not available. The dietary uptake formulation represented the metabolite-level average intake of a one-day meal plan for a 70 kg adult^28^ and was added to the models using the *setDietConstraints.m* CobraToolbox function. The applied diet contained 192 metabolites (Table S3), of which 92 metabolites were present in trace amounts below 0.1 mmol/day/person.

### 4.10 Simulations

The metabolic fluxes in blood were predicted by performing FBA, which converts a genome-scale metabolic reconstruction into a mathematical model and interrogates the model using a linear programming solver according to an objective function^15^. The objective function describes a linear optimisation problem for finding extreme points in a constrained solution space of metabolic states as follows:

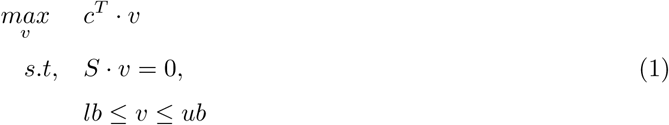

The linear programming solver then finds a reaction flux vector, *v*, containing a maximised metabolic flux through an objective reaction, defined by a non-zero coefficient in *c*. The maximised flux value must lie within the feasible solution space defined by constraints, such as the steady state constraint. Steady state is defined by the system of linear equations, *S · v* = 0, which enforces unchanging metabolite concentrations over time. Consequently, the amount of produced metabolites (products) equalled the amount of consumed metabolites in each reaction (substrates). The metabolic products and substrates in each reaction are encoded by non-zero coefficients in the stoichiometric matrix, *S*. Substrates are represented by a negative coefficient, while reaction products are indicated by positive coefficients. In addition to the steady state constraint, each reaction in *v* was constrained by reaction-specific upper (*ub*) and lower (*lb*) bounds, derived from literature sources such as thermodynamic data and experimental measurements. If no reaction information was available, reactions were left unconstrained by setting *lb* and *ub* to arbitrary values of -1,000,000 and 1,000,000 mmol/person/day (see also Thiele et al.^21^ for details). For the objective reactions, we introduced demand reactions into the WBM blood compartments (VMH name: [bc]) for each of the analysed blood metabolites. Demand reactions are unbalanced, left-sided reactions (e.g., DM leu L[bc]: 1 leu L[bc] *→* Ø for L-leucine in blood), commonly used to predict the maximal potential metabolite productions in a system. In previous studies, demand reactions have been used in WBMs to predict blood biomarkers of single gene defects^112^ and gut microbiome flux influences associated with risk markers of Alzheimer’s disease^24^. Demand reactions were separately introduced for each FBA using the *addDemandReaction.m* CobraToolbox function^107^. FBA was then performed for each model by running *fba = optimizeWBModel(model)* with the *model.osenseStr = ’max’* parameter to ensure maximisation of the flux through the objective reaction. Default parameters were used for the linear optimisations. After performing FBA, the maximised fluxes were extracted from the *fba.f* solution field and rounded to six decimals to remove potential numerical artefacts that can appear in small numbers.

### 4.11 Outlier removal

Outlier samples in the predicted fluxes were identified using a robust principal component analysis (PCA) on the z-transformed fluxes, performed with the alternating least squares algorithm. Potential outliers were identified by taking the Euclidean norm of the principal component scores for each sample, weighted by the explained variance of each principal component. This approach obtains the average distance of a sample in a principal component from the centre of the flux data. Values close to zero indicate samples for which average fluxes were predicted, while the highest calculated values in the cohort were flagged as potential outlier samples across the predicted fluxes. In combination with a visual inspection of the first three PCA components, the ranked outlier scores were used to manually identify outlier samples to be removed.

### 4.12 Statistical analyses

#### 4.12.1 Study cohort descriptions

The Fisher’s exact test was used to quantify sample differences in AD patients and controls for the binary variables in the metadata. The *fishertest.m* function was used to calculate the Fisher-exact p-values, the associated odds ratios, and the 95% confidence intervals. For the numerical metadata, including age in years and the mapped read count, differences between AD patients and controls were tested by performing the Wilcoxon rank sum test, using the *ranksum.m* MATLAB function.

#### 4.12.2 AD dementia analysis

Multiple linear regressions were performed to identify differences in the predicted blood flux values (response) with *AD* status (predictor). The linear regressions were stratified for pairwise investigations on each AD status group comparison, i.e., healthy control versus AD MCI, AD MCI versus AD dementia, and healthy control versus AD dementia. The linear regressions further included sex and age at stool sample collection (Z-transformed) as household control variables. Furthermore, technical control variables were added for the sequencing lane, whether ethanol was added during the DNA extraction (yes/no), and the total reads in each microbiome-personalised WBM (log10-transformed and Z-transformed). The same set of multiple linear regressions was performed to associate the gut metagenomics reads in the microbiome-personalised WBMs and the plasma metabolomic abundances (log-transformed and Z-transformed). These regressions did not include control variables for the microbiome-specific covariates, including the sequencing lane, whether ethanol was added during the DNA extraction (yes/no), and the total reads in each microbiome-personalised WBM.

#### 4.12.3 Flux to plasma metabolomic association analysis

To assess the associations between predicted blood fluxes and the corresponding plasma metabolomics, linear regressions were performed on the log2-transformed and z-scaled flux predictions (predictor) against the z-scaled and log-transformed metabolomic plasma abundances (response) with control variables for sex, age, the total reads in each microbiome-personalised WBM (log10-transformed and Z-transformed), and the sequencing lane.

#### 4.12.4 *APOE* risk group analysis

Multiple linear regressions were performed to identify differences in the predicted blood flux values (response variable) across *APOE* risk group status (predictor). The linear regressions were stratified by pairwise investigations on each *APOE* risk group comparison, i.e., *ε*2 versus *ε*3, *ε*3 versus *ε*4, and *ε*2 versus *ε*4. Each linear regression included sex and age at stool sample collection (Z-transformed) as household control variables. Furthermore, technical control variables were added for the sequencing lane, if ethanol was added during the DNA extraction (yes/no), and the total reads in each microbiome-personalised WBM (log10-transformed and Z-transformed). The same set of multiple linear regressions was performed to investigate associations between the gut metagenomics reads in the microbiome-personalised WBMs and the plasma metabolomic abundances (log-transformed and Z-transformed). However, the regressions on the plasma metabolomics did not include control variables for the microbiome-specific covariates, such as the sequencing lane, whether ethanol was added during the DNA extraction (yes/no), and the total reads in each microbiome-personalised WBM.

#### 4.12.5 MoCA score analysis

Multiple linear regressions were performed on the Z-scaled MoCA scores (response) with the predicted blood flux values as the predictor of interest. These linear regressions included age and sex as household covariates and technical covariates for the sequencing lane, if ethanol was added during the DNA extraction (yes/no), and the total reads in each microbiome-personalised WBM (log10-transformed and Z-transformed). Additional control variables included BMI in kg / *m*^2^ (log10-transformed and Z-transformed), the number of years of education, and *APOE ε*4 carrier status (yes/no) were added as control variables. The same set of multiple linear regressions was performed to investigate the gut metagenomics reads in the microbiome-personalised WBMs and the plasma metabolomic abundances (log-transformed and Z-transformed). However, the regressions on the plasma metabolomics did not include control variables for the microbiome-specific covariates, including the sequencing lane, whether ethanol was added during the DNA extraction (yes/no), and the total reads in each microbiome-personalised WBM.

### 4.13 Identification of key microbes for the flux predictions

#### 4.13.1 Shadow price analysis

Potential microbial influencers of the predicted fluxes were identified by analysing the shadow prices in the FBA solutions associated with the flux predictions. Shadow prices are solutions to the dual linear problem and represent the sensitivity of the flux prediction upon a change in metabolite availability. More precisely, the shadow price of a metabolite *i*, *π_i_*, represents the inverse ratio of an incremental relaxation of the steady state constraint for a metabolite *i*, *δb_i_*, and the resulting maximal change in the flux through the objective reaction, *δZ*:

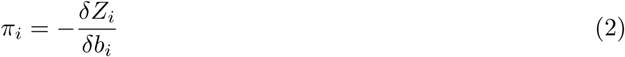

To identify the potential microbial influencers of the flux predictions, we analysed the shadow prices of the microbial biomass metabolites in the WBMs. Microbial biomass metabolites were coupled with the metabolic reactions in the microbial pan model via flux capacity (coupling) constraints^110^, which enforced a minimal flux through the microbial reactions at a fixed ratio of 400 times the flux through the biomass reaction. A consequence of these biomass reaction couplings is that if a microbial species produces any metabolite in the pathway of the predicted metabolites in blood, a non-zero shadow price is found. Therefore, the shadow prices of the microbial biomass metabolites reveal which microbial species can contribute to the predicted fluxes. These shadow prices were automatically computed when performing FBA and were extracted from *fba.y* after performing *fba = optimizeWBModel(model)*. Potential microbial influencers were then identified for each FBA prediction by assessing if the shadow price value was zero, meaning that a change in microbial biomass would not change the flux prediction, or non-zero, indicating that altered microbial abundances could influence the flux results. A numerical tolerance for the shadow prices of 1e-6 was used to remove non-zero shadow prices resulting from numerical imprecision. After extracting the microbial species with non-zero shadow prices in each FBA prediction, microbial species with an average non-zero biomass shadow price for their associated flux predictions across all analysed samples were identified by calculating the bootstrapped average and associated 95% confidence interval for the microbial biomass shadow price across the analysed samples and each metabolic blood flux prediction. Bootstrapping was performed with 10,000 samples with replacement. Microbial species with cohort-average shadow prices that did not include zero in their 95% confidence intervals were flagged as potential contributors to the altered flux predictions in AD patients. Taken together, potential microbial influencers of the flux predictions were identified by analysing the flux-associated shadow prices of the pan-biomass metabolites of microbial species included in the microbiome-personalised WBMs.

#### 4.13.2 Selection of flux-associated microbes

Flux-associated microbial species with potential relevance to the altered flux predictions in AD patients were identified using the stability selection method as described by Meinshausen and Bühlmann^35^. This method was chosen as it 1) identified relevant variables in higher-dimensional datasets, 2) accounted for confounding influences on the fluxes as a result of the sex-specific metabolic structure of the WBMs, and 3) provided stable variable selection outcomes across function calls. The stability selection method relies on LASSO regressions, which extend the linear regression paradigm by adding a term that minimises the L1 norm of the fitted regression coefficients. Consequently, fitted LASSO regression coefficients reflect the minimal sum of squared errors for the smallest of predictors with non-zero coefficients. The weight of the additional term in LASSO regression was defined by a regularisation parameter *λ*. LASSO regressions were performed using the lasso.m function, which automatically selects the optimal *λ* by identifying a global optimum for the LASSO objective function using the coordinate descent algorithm described by Friedman et al.^113^. Stability selection was performed by creating 500 random samples, each containing 70% of the analysed data and performing LASSO regressions with predictors for the sex variable and the z-scaled relative abundances of the flux-associated microbial species. The z-scaled predicted flux was used as the response variable in the regressions. For each of the 500 samples, the microbial species with non-zero regression coefficients were extracted, and their selection frequency was counted. Microbial species were selected for further analysis if they had a non-zero regression coefficient in at least 90% of the 500 samples. To summarise, the stability selection method was used to identify microbial species that improved the regression fit for associating microbial relative abundances with the AD-associated blood fluxes.

#### 4.13.3 Calculation of microbe importances for the predicted blood fluxes

Microbial importances for the flux predictions were obtained by performing elastic net regressions^37^ on the z-scaled flux predictions with predictors for the relative abundances of the selected microbial species and the sex variable. The elastic net regressions were performed using the lasso.m function with a *α* weight parameter of 0.5, which assigns an equal weight to the LASSO and ridge term of the elastic net objective function. The regularisation parameter *λ* was estimated by performing ten-fold cross-validation and identifying the *λ* value corresponding to the smallest mean square error of the regression fit.

### 4.14 Software

Modelling work, including the construction and analysis of the microbiome WBMs, was done in MATLAB 2020b and MATLAB 2024b (Mathworks^TM^). In addition, the parallel computing toolbox, statistics and machine learning toolbox, and the bioinformatics toolbox were utilised to generate and analyse microbiome WBMs. The generation and analysis of the microbiome-personalised WBMs were performed using functions from the COBRA toolbox v3^107^, including the microbiome modelling^108^ and PSCM^21^ toolboxes. The IBM ILOG CPLEX 12.10 linear solver (IBM Inc.) was used for performing FBA on the investigated metabolites. Visualisation was performed in MATLAB and R version 4.5.0 using the tidyverse family of packages^114^ and the pheatmap package^115^.

## 5 Code availability

All scripts can be accessed at https://github.com/ThieleLab/CodeBase.

## 6 Supplemental material

Supplementary tables S1–S9 can be accessed at: https://docs.google.com/spreadsheets/d/1fJXCbaUBCK569zswkLuSqL-FGlroQeDj/edit?usp=sharing&ouid=108109531682123793728&rtpof=true&sd=true

## 7 Data availability statement

Samples were provided to NCRAD by ten Alzheimer’s Disease Research Centres, including UC San Diego, UC Davis, Stanford University, Kansas University, Wisconsin University, Indiana University, New York University, Wake Forest University, Cleveland Clinic, Nevada, and the University of Alabama at Birmingham. Clinical data can be requested from the National Alzheimer’s Coordinating Centre (naccdata.org/). Data will be available in the Synapse AD Knowledge Portal. Gut microbiome data are stored and accessible via the University of California, San Diego Qiita platform (qiita.ucsd.edu/). The AGORA2 reconstructions can be freely downloaded from the VMH resource (https://www.vmh.life/files/reconstructions/AGORA2/), while the APOLLO reconstructions are available from the Harvard Dataverse (version 1.0 https://doi.org/10.7910/DVN/PIZCBI).

## 8 Acknowledgments

This project was enabled in part by the AGMP and Alzheimer Disease Metabolomics Consortium (ADMC), supported by the National Institute on Aging grants: R01AG046171, RF1AG051550, RF1AG057452, R01AG059093, U19AG063744, 3U19AG063744-04S1, RF1AG058942, 1U01AG088562, U01AG061359, R01MH108348, R01AG081322 and FNIH: #DAOU16AMPA, awarded to Dr. Kaddurah-Daouk at Duke University in partnership with multiple academic institutions. As such, the investigators within the AGMP not listed in this publication’s authors’ list, provided analysis-ready data, but did not participate in designing the study, conducting the analyses or writing of this manuscript. A listing of AGMP Investigators can be found at https://alzheimergut.org/meet-the-team/. A complete listing of the AD Metabolomics Consortium (ADMC) investigators can be found at: https://sites.duke.edu/adnimetab/team/. The NACC database is funded by NIA/NIH Grant U24 AG072122. NACC data are contributed by the NIA-funded ADRCs: P30 AG062429 (PI James Brewer, MD, PhD), P30 AG066512 (PI Thomas Wisniewski, MD), P30 AG072976 (PI Andrew Saykin, PsyD), P30 AG072975 (PI Julie A. Schneider, MD, MS), P30 AG062715 (PI Sanjay Asthana, MD, FRCP), P30 AG072973 (PI Russell Swerdlow, MD), P30 AG072947 (PI Suzanne Craft, PhD), and P30 AG086401 (PI Erik Roberson, MD, PhD). P30 AG072972 (PI Charles DeCarli) and P30 AG066515 (PI Victor Henderson) contributed microbiome samples to AGMP via NCRAD, but not plasma. The IGM S10 grant S10 OD026929 was awarded to Dr. Rob Knight at the University of California, San Diego, IGM Genomics Centre. This publication includes data generated at the University of California, San Diego IGM Genomics Centre utilising an Illumina NovaSeq 6000 that was purchased with funding from an NIH SIG grant (S10 OD026929). Lucas Patel is supported by the University of California, San Diego Medical Scientist Training Program (NIH/NIGMS T32GM007198). Samples from the National Centralised Repository for Alzheimer’s Disease and Related Dementias (NCRAD), which receives government support under a cooperative agreement grant (U24 AG021886) awarded by the National Institute on Aging (NIA), were used in this study. We thank contributors who collected samples used in this study, as well as patients and their families, whose help and participation made this work possible. Further funding for this project was provided through the European Union’s Horizon 2020 research and innovation programme [grant agreement No 757922 and 101125633] to Dr. Ines Thiele and from the Science Foundation Ireland under [Grant number 12/RC/2273-P2]. This research was further funded by the Health Research Board (JPND-2023-3). The authors would also like to thank and acknowledge Paula Walsh, Naveen James, and Dr. Cyrille Thinnes for their professional support and valuable advice and Dr. Naama Karu for her valuable feedback on the paper. The authors would further like to thank and acknowledge Dr. Hazel Dilmore and Dr. Mackenzie Bryant for their contributions to the curation, processing, and management of the gut metagenomics data.

## 9 Conflict of interest statement

Dr. Kaddurah-Daouk is an inventor of a series of patents on the use of metabolomics for the diagnosis and treatment of central nervous system (CNS) diseases and holds equity in Metabolon Inc. and Chymia LLC. Dr. Rob Knight is a scientific advisory board member and consultant for BiomeSense, Inc., has equity, and receives income. He is a scientific advisory board member and has equity in GenCirq. He is a consultant and scientific advisory board member for DayTwo and receives income. He has equity in and acts as a consultant for Cybele. He is a co-founder of Biota, Inc., and has equity. He is a co-founder of Micronoma and has equity and is a scientific advisory board member. The terms of these arrangements have been reviewed and approved by the University of California, San Diego, in accordance with its conflict of interest policies. Daniel McDonald is a consultant for BiomeSense, Inc., has equity, and receives income. The terms of these arrangements have been reviewed and approved by the University of California, San Diego, in accordance with its conflict of interest policies. All other authors have no conflicts of interest to disclose beyond those listed.

## 10 Consent statement

All human subjects provided written informed consent to participate in this study.

## 11 Author Contributions

CRediT: Tim Hensen: Conceptualisation, Formal analysis, Investigation, Methodology, Visualisation, Writing – original draft, Writing – review & editing; Lora Khatib: Data curation, Formal analysis; Lucas Patel: Data curation, Formal analysis; Daniel McDonald: Data curation, Formal analysis; Antonio González: Data curation, Formal analysis; Siamak MahmoudianDehkordi: Data curation, Formal analysis; Colette Blach: Project administration; Rob Knight: Funding acquisition, Resources, Supervision, Writing – review & editing; Rima Kaddurah-Daouk: Funding acquisition, Resources, Supervision, Writing – review & editing; Ines Thiele: Conceptualisation, Methodology, Funding acquisition, Resources, Supervision, Writing – review & editing.

